# TET3 regulates cellular terminal differentiation at the metabolic level

**DOI:** 10.1101/2024.01.23.576868

**Authors:** Isabel Mulet, Carmen Grueso-Cortina, Mireia Cortés-Cano, Daniela Gerovska, Guangming Wu, Daniel Jimenez-Blasco, Andrea Curtabbi, Pablo Hernansanz-Agustín, Israel Manjarrés-Raza, Juan Pedro Bolaños, José Antonio Enríquez, Marcos J. Araúzo-Bravo, Natalia Tapia

## Abstract

TET-family members play an essential role in cell fate commitment and their dysfunctions result in arrested differentiation. TET3 is ubiquitously expressed in differentiated cells and essential in postnatal development due to yet unknown reasons. To define TET3 function in cell differentiation, we profiled the intestinal epithelium at the single-cell level from wild-type and *Tet3* knockout mice. Here we show that, in the absence of TET3, enterocytes exhibit an aberrant differentiation trajectory and do not acquire a physiological cell identity due to an impairment in oxidative phosphorylation, specifically due to an ATP synthase assembly deficiency. Furthermore, our analysis demonstrates that the loss of TET3 compromises mitochondrial metabolic maturation and leads to a metabolic profile enriched in glycolysis-dependent anabolic pathways similar to those observed in undifferentiated cells. Collectively, our study has revealed the molecular mechanism by which TET3 regulates terminal differentiation at the metabolic level.

## INTRODUCTION

DNA methylation of the cytosine base at the fifth carbon position (5-methylcytosine; 5mC) is a key epigenetic mechanism involved in development, genome regulation and disease^1^. In mammalian cells, 5mC can be oxidized into 5-hydroxymethylcytosine (5hmC)^2,3^. To date, the *Ten-eleven-translocation* gene family (*Tet1*, *Tet2* and *Tet3*) are the only enzymes that have been described to convert 5mC into 5hmC through their α-ketoglutarate/Fe(II)-dependent dioxygenase activity^4^.

5hmC associates with the activation of key genes during progenitor differentiation, suggesting a general role of 5hmC in establishing cell identity^4^. In the intestinal epithelium, 5hmC globally accumulates in the intestinal stem cell progeny as they mature^5^. In the ectodermal lineage, global 5hmC levels increase during neural differentiation and double*Tet2* and *Tet3* depletion in the mouse cortex blocks progenitor differentiation into neurons^6^. In contrast, hematopoietic stem cell differentiation progresses with a global decrease in 5hmC. Despite this genome-wide 5hmC reduction, 5hmC enrichment at particular loci is required for hematopoietic progenitors to properly differentiate and an inability to do so leads to a biased differentiation into the myeloid lineage^4^. Overall, the TET enzymes seem to play an essential role on lineage differentiation in all three germ layers and, accordingly, TET dysfunctions result in skewed or arrested differentiation.

Loss-of-function mouse models from the three *Tet* family members have provided insight into their function and redundancy. *Tet1* knockout mice are embryonically lethal in outbred mice but viable in an inbred strain and the reason for this phenotypic discrepancy remains controversial^7,8^. On the contrary, *Tet2*-null mutants survive to an adult stage but develop lethal myeloid malignancies^9^. Double *Tet1* and *Tet2* deficient mice are born at mendelian ratios but half of them die perinatally, indicating that *Tet1* and *Tet2* enhance survival but that their loss does not impair postnatal development^10^. Remarkably, *Tet3* knockout mice develop to term but die perinatally due to unknown reasons, suggesting that *Tet1* and *Tet2* cannot compensate for *Tet3* critical functions at birth^11^. Overall, the *Tet* paralogs exhibit functional differences that are currently not well-understood.

TET3 deficiency provokes developmental defects in multiple cell types indicating that TET3 is a crucial factor for cellular commitment and terminal differentiation^12^. In adult homeostasis, the role of *Tet3* has been studied in a few tissues together with *Tet2*, as both paralogs are ubiquitously expressed in most differentiated cells^4,13^. Indeed, TET3 can rescue TET2 loss during hematopoietic differentiation to some lineages but not to others, suggesting that TET2 and TET3 share some but not all targets^4^. Mechanistically, TET3 appears to be recruited to specific gene loci by transcription factors since TET3 has no DNA-binding specificity itself^12^. Indeed, TET3 has been described to be recruited to certain lineage-associated genomic loci via interaction with FOXA2 and C/EBPδ^12,14^, to catalyze DNA demethylation and stimulate the expression of key genes, thus controlling cell differentiation. It has also been hypothesized that an increase in DNA hydroxymethylation levels might enhance chromatin accessibility and binding of key lineage specific transcription factors^13^. However, the precise molecular mechanism in which TET3 physiologically regulates adult stem cell differentiation remains practically unresolved.

In this study, we have used the mouse intestinal epithelium as a model to investigate the role of TET3 during adult stem cell differentiation into their progeny. Our data demonstrates that TET3 is required for ATP synthase assembly and, thus, for mitochondrial maturation during cell fate commitment. Therefore, in the absence of TET3, enterocytes do not acquire a mature cell identity due to a deregulation in oxidative phosphorylation.

## RESULTS

### Global 5hmC accumulation during enterocyte maturation is mainly TET3-dependent

To better understand the precise physiological role of TET3 in stem cell-hierarchical tissues, we generated a heterozygous *Tet3* knockout mouse line through homologous recombination in which exon 5 was excised (Supp. Fig. 1a, d). Homozygous knockout mice were born at predicted mendelian ratios but died on postnatal day 1 showing no gross abnormalities (Supp. Fig. 1b, c). 5-hydroxymethylcytosine (5hmC) staining on whole E18.5 embryo sections pointed to the intestinal epithelium as the tissue with the most significant reduction in 5hmC levels in the absence of TET3 (data not shown), indicating that TET3 is the *Tet*-enzyme responsible for maintaining 5hmC physiological levels in the intestinal epithelium. Hence, we decided to focus on the intestinal epithelium for further studies since we assumed that the substantial decrease of 5hmC observed in this tissue might help us to pinpoint TET3 critical functions. In the wild-type, intestinal stem cells are 5hmC—negative in contrast to their progeny, in which 5hmC globally accumulates as they mature and progress upwards through the intervilli-villi axis (Fig. 1a). Interestingly, the *Tet3* knockout intestinal epithelium exhibited a reduction in the intensity and number of 5hmC—positive cells whereas the 5hmC levels on lamina propria and interstitial space were unaffected (Fig. 1a). Indeed, liquid chromatography-tandem mass spectrometry showed a 50% global genome 5hmC decrease in the *Tet3* knockout intestinal epithelium (Fig. 1b). On the contrary, 5-methylcytosine levels remain unchanged after TET3 depletion, excluding TET3-mediated 5hmC deposition in the intestinal epithelium as a mere intermediate of DNA demethylation (Fig. 1a,b). Quantitative RT-PCR on intestinal epithelial cells demonstrated that *Tet2* and *Tet3* are expressed at similar levels whereas *Tet1* is almost undetectable. Remarkably, *Tet3* expression silencing was accompanied by a slight but significant increase in *Tet2* expression in the *Tet3* knockout (Fig. 1c). To investigate whether all the intestinal stem cell progeny or whether only specific daughter cell types showed a decrease in 5hmC signal, we performed a double immunostaining with 5hmC together with cell-type specific markers. The quantification showed that the number of 5hmC—positive enteroendocrine and goblet cells is unaltered in the absence of *Tet3* in contrast to the number of 5hmC—positive enterocytes that is 70% reduced. Intestinal stem cells that are located in the intervillus region are 5hmC— negative, as expected (Fig. 1d). Collectively, our results demonstrate that *Tet3* is the main *Tet*-family member responsible for 5hmC deposition in enterocytes.

**Fig. 1.**
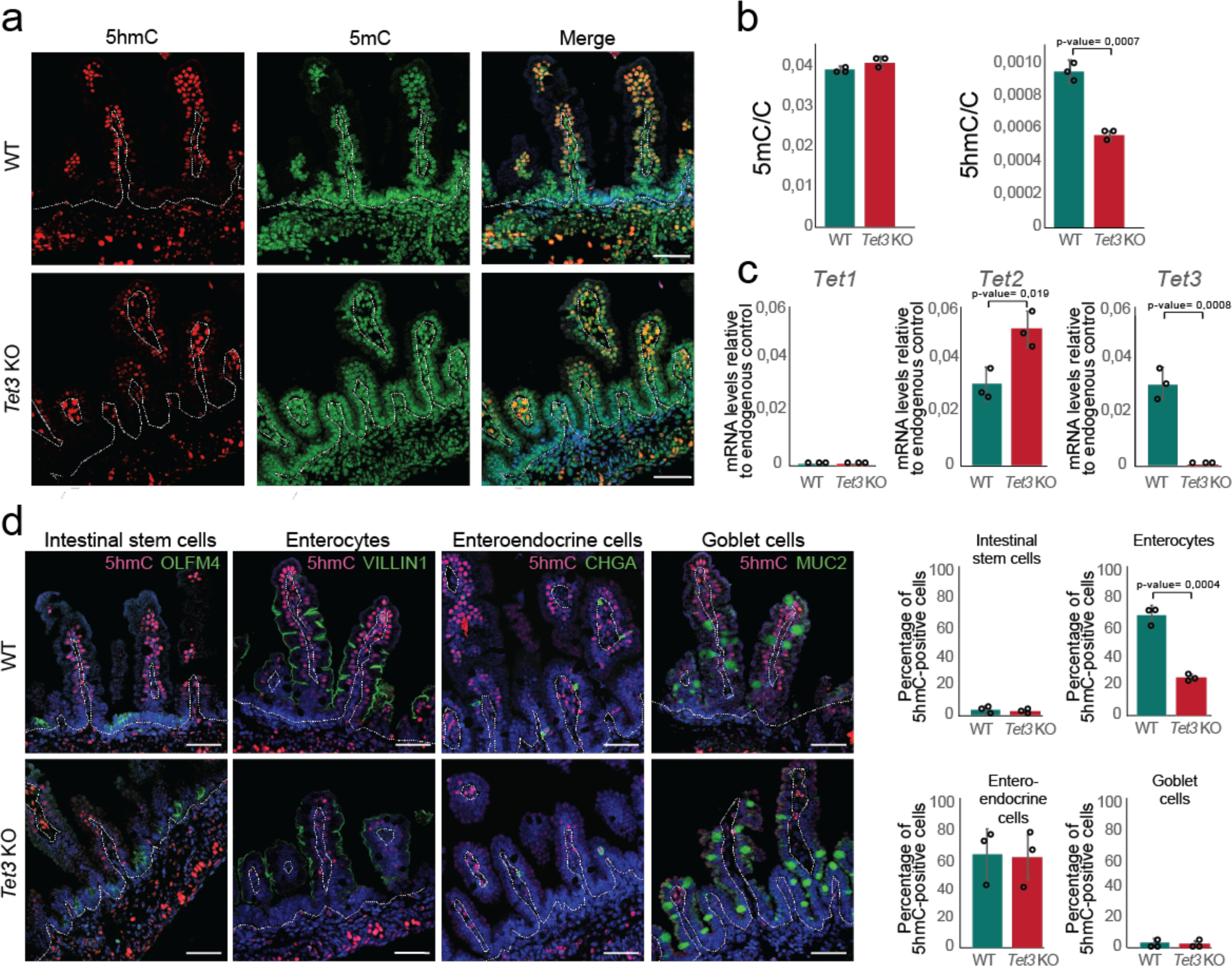
Global 5hmC accumulation during enterocyte maturation is mainly TET3-dependent. **a** Immunofluorescence analysis of 5hmC and 5mC in E18.5 small intestines. Nuclei were stained with DAPI. **b** Mass spectrometry quantification of 5mC and 5hmC relative to C in E18.5 intestinal epithelial cells. Data are mean ± s.d. (n=3 biologically independent samples). **c** *Tet* expression levels in E18.5 intestinal epithelial cells were measured by RT-qPCR and represented as relative to the endogenous control. Each sample corresponds to a pool of three E18.5 intestinal epitheliums. Error bars represent standard deviation of technical replicates. **d** Double immunofluorescence quantification of 5hmC and intestinal cell type-specific markers in E18.5 small intestine. Nuclei were stained with DAPI. Data are mean ± s.d. (n=3 biologically independent samples). Dashed line delineates intestinal epithelium. Scale bars represent 50μm. C, total cytosines; 5mC, 5-methylcytosines; 5hmC, 5-hydroxymethylcytosines. Student’s *t* test, two sided.

### TET3 is required to acquire a physiological enterocyte signature

Next, we investigated the role of 5hmC on enterocyte transcriptional regulation. To this end, we used the single cell 10X Genomics platform to profile single cell (sc) RNA-seq of 12,000 individual EpCam—positive epithelial cells sorted from the small intestine of wild-type and *Tet3* knockout E18.5 embryos (Supp. Fig. 2a). Our analysis resolved seventeen clusters that were associated with distinct cell identities namely, stem/transit-amplifying cells (clusters 4, 9, 11, 12 and 14), goblet cells (cluster 16), proximal enterocyte (clusters 1, 2, 3, 5, 6, 7, 8, 10 and 13) and distal enterocyte (cluster 0), based on known markers from adult intestinal epithelium, since the information available for newborns is limited (Fig. 2a and Supp. Table 1). The top 100 genes defining each cluster are shown in Supp. Table 2. Both wild-type and *Tet3* knockout goblet cells are represented by a single distinct cluster (cluster 16) that contains a similar percentage of wild-type and knockout cells (Supp. Table 3). Likewise, no major differences in the number of cells of each genotype are observed in clusters 9, 11, 12, 4 and 14 that correspond to stem/transit-amplifying cells (Supp. Table 3). These results are consistent with the immunostaining analysis (Fig. 1). In contrast, enterocytes segregate in different clusters depending on their genotype, in which clusters 0, 1, 5 and 7 are mainly composed of wild-type cells whereas clusters 2, 3, 8 and 13 are composed almost 99% of *Tet3* depleted cells (Fig. 2b, c and Supp. Table 3). A trajectory analysis confirmed a bifurcating trajectory from stem/transit-amplifying cells into the secretory and the absorptive lineage and a totally different enterocyte differentiation trajectory between the wild-type and the *Tet3* knockout cells (Fig. 2d). Indeed, the wild-type cells progress from the stem/transit-amplifying cells into enterocytes (clusters 0, 1, 5 and 7). Instead, the majority of the *Tet3* knockout cells follow an aberrant trajectory and accumulate in clusters 2, 3, 8 and 13, indicating that most cells do not reach a full mature enterocyte identity in the absence of TET3.

**Fig. 2.**
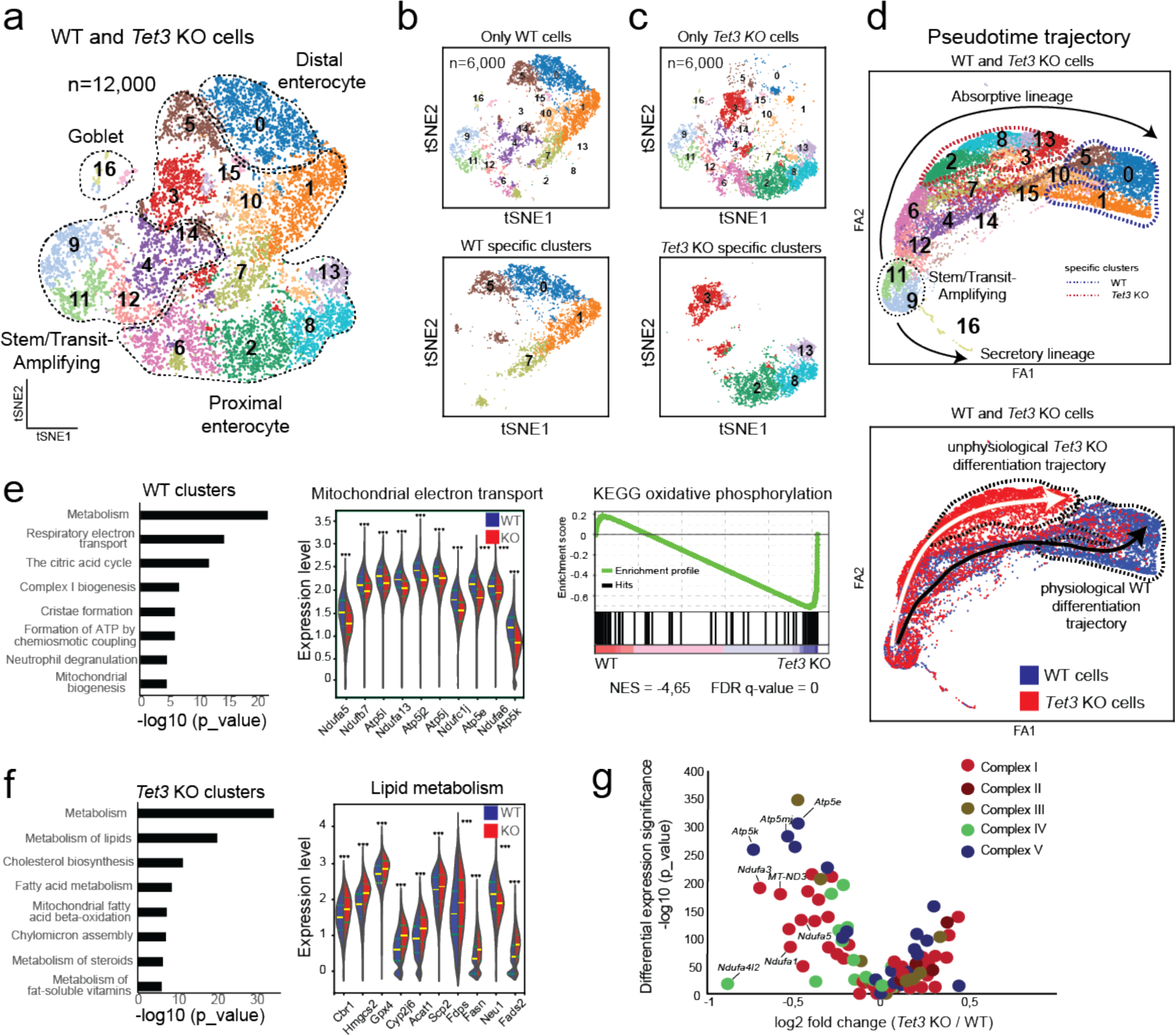
Enterocyte acquire an aberrant transcriptional signature in the absence of TET3. **a** Cluster color-coded scRNA-seq t-SNE visualization of 6,000 E18.5 wild-type and 6,000 E18.5 *Tet3* knockout small intestine epithelial cells. Clusters are labelled with numbers and cell identities are delineated with dashed lines. **b** t-SNE visualizations of (upper panel) only wild-type cells n=6,000 cells and (lower panel) only wild-type specific clusters n=3,816 cells. **c** t-SNE visualizations of (upper panel) only *Tet3* knockout cells n=6,000 cells and (lower panel) only *Tet3* knockout specific clusters n=3,211 cells. **d** Cluster color-coded (upper panel) or genotype color-coded (lower panel) cell differentiation trajectory containing both wild-type and *Tet3* knockout cells. n=6,000 cells for each genotype. Black and white arrows in lower panel show differentiation into the physiological and aberrant absorptive lineage, respectively. Dashed lines delimit wild-type and *Tet3* knockout specific clusters. **e**, **f** (left panel) **e** Top 8 significantly enriched ontology terms identified using g:Profiler in specific wild-type clusters (n=3,816 cells) and **f** *Tet3* knockout specific clusters (n=3,211 cells), respectively. **e**, **f** (middle panel) **e** Expression levels of genes representative of the mitochondrial electron transport pathway or **f** lipid metabolism pathway in wild-type (n=3,816 cells) and *Tet3* knockout specific clusters (n=3,211 cells). Median as well as the first quartile and the third quartile are depicted in green and yellow, respectively. ***p<0.001. **e** (right panel) GSEA plot showing oxidative phosphorylation as a negatively enriched pathway in *Tet3* knockout epithelial cells. **g** Differential expression and statistical significance between wild-type (n=3,816 cells) and *Tet3* knockout (n=3,211 cells) specific clusters for mitochondrial subunits showing p-values < 0,05.

A gene ontology analysis was performed to identify the main biological functions enriched in wild-type (clusters 0, 1, 5 and 7) and in aberrant (clusters 2, 3, 8 and 13) enterocytes (Fig. 2e). The analysis revealed a significant enrichment of oxidative phosphorylation in the wild-type but of fatty acid metabolism and sterol biosynthesis in the *Tet3* knockout enterocytes (Fig. 2f), suggesting that energy metabolism is dysregulated in the absence of TET3. Indeed, a volcano plot of all the components of the mitochondrial electron transport chain showed that a large number of complex I and F_1_F_0_ATP synthase—complex V subunits were significantly downregulated in the *Tet3* knockout enterocytes (Fig. 2g). Overall, our results demonstrate that the loss of TET3 leads to enterocytes with an unphysiological metabolic transcriptional signature.

### Mitochondrial morphology is abnormal in the absence of TET3

The gene expression analysis suggests a dysregulation in oxidative phosphorylation in the absence of TET3 (Fig. 2g). Next, we sought to ascertain the impact of TET3 loss on mitochondrial morphology as it associates with mitochondrial function^15^. In the intestinal epithelium, while the intestinal stem cells rely largely on glycolysis to fulfill their energetic demands, enterocyte dependence on oxidative phosphorylation increases with differentiation^16^. Mitochondrial elongation has been suggested to correlate with oxidative metabolism^17^ and, accordingly, enterocytes exhibit more elongated mitochondria than intestinal stem cells^18^. Hence, electron microscopy analysis was employed to study the mitochondria size in the villi tip and in the intervilli region in both wild-type and *Tet3* knockout (Fig. 3a). As previously reported, the wild-type mitochondria are significantly larger in the differentiated cells located at the villus tip than at the stem cell compartment (Fig. 3b). In contrast, the mitochondrial area does not increase in the *Tet3* knockout villi tip, pointing to a failure in mitochondria maturation during intestinal epithelial cell differentiation in the absence of TET3 (Fig. 3b). Interestingly, there is no difference in mitochondrial area between the wild-type and the *Tet3* knockout mitochondria located in the intervilli (Fig. 3b). These findings suggest that TET3 loss has no impact on the stem cell compartment and correlate with the results obtained by immunostaining (Fig. 1d) and global gene expression profiling (Fig. 2d). In contrast, mitochondrial size is significantly larger in the wild-type in comparison with the *Tet3* knockout at the villi tip (Fig. 3b), indicating that mitochondrial maturation does not properly progress upon TET3 loss. Overall, mitochondria show an abnormal morphology in differentiated cells in the absence of TET3, which points to an alteration in mitochondrial fate during intestinal epithelial differentiation.

**Fig. 3.**
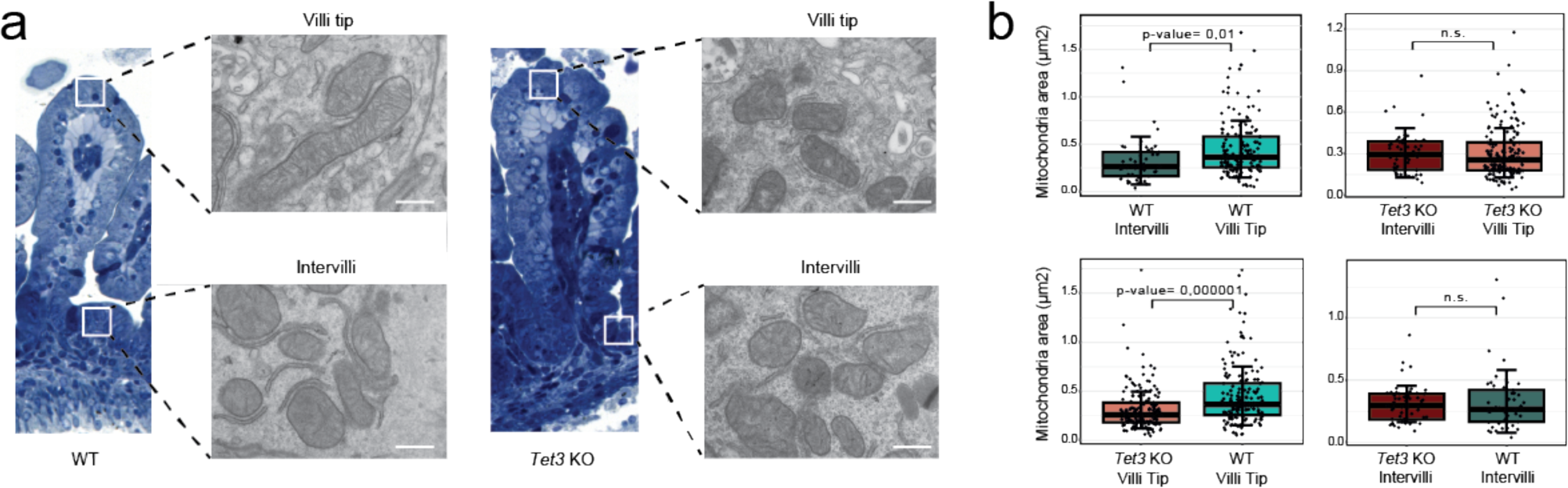
Mitochondrial morphology remains immature in the absence of TET3. **a** Representative transmission electron microscopy images from E18.5 wild-type and *Tet3* knockout mitochondria located at intervilli and villi tips are shown. Semi-thin sections images with boxed areas indicating the analyzed villi regions are included. Scale bars: 500 nm. **b** Quantitative mitochondrial area comparison between wild-type intervilli, *Tet3* knockout intervilli, wild-type villi tip and *Tet3* knockout villi tip (n=45, 53, 155, and 150 mitochondria, respectively). Error bars correspond to the standard deviation. Student *t* test, two-tailed.

### TET3 regulates mitochondrial F_1_F_0_-ATP synthase assembly

*Tet3* knockout enterocytes showed a reduction in the expression of a large number of mitochondrial complex I and complex V subunits, thus we next attempt to functionally validate these results. However, isolating wild-type (clusters 0, 1, 5 and 7) and *Tet3* knockout (clusters 2, 3, 8 and 13) differentiated enterocytes is technically not feasible. For this reason, we decided to assess mitochondrial complex I and complex V activities in E18.5 tail tip derived-fibroblasts that, like enterocytes, also exhibit a terminally differentiated phenotype. Indeed, quantitative gene expression analysis in *Tet3* knockout fibroblasts revealed a decrease in the expression of several mitochondrial subunits at levels similar to those observed in *Tet3* knockout enterocytes (Supp. Fig. 3a). Differences in complex I assembly were not observed despite the reduction in gene expression of many of its subunits, as shown by blue native gel electrophoresis (BNGE) followed by either in gel activity or western blotting for NDUFA9 (Fig. 4a). Next, we assessed the function of mitochondrial complex I that is known to transition between active and deactive forms^19^. NADH:ubiquinone activity informs on the ability of complex I to extract electrons from NADH and transfer them to ubiquinone, allowing to determine the active form. On the other hand, NADH:Fe(CN)_6_ oxidoreductase activity informs on the available complex I regardless if it is in active or deactive form. For the purpose to estimate both the amount of complex I and the proportion of active complex I, we measured NADH:ubiquinone and NADH:Fe(CN)_6_ oxidoreductase activities spectrophotometrically in isolated mitochondria from wild-type and *Tet3* knockout (Fig. 4b). Our results show a significant decrease in complex I NADH:Fe(CN)_6_ but not in NADH:ubiquinone oxidoreductase activity, suggesting that the percentage of activated complex I is higher in *Tet3* knockout fibroblasts and compensates for the reduction in the total amount of complex I in the absence of TET3. Self-association of F_1_F_0_-ATP synthase monomers into rows of dimers and oligomers is a fundamental biological process involved in energy efficiency^20^. In mammals, F_1_F_0_-ATP synthase dimerization/oligomerization is mediated by specific subunits belonging to the F_1_F_0_-ATP synthase membrane domain. Strikingly, most of the F_1_F_0_-ATP synthase subunits that were downregulated in the *Tet3* knockout fibroblasts are located in the membrane domain (Fig. 4c). Next, we proceed to use BNGE to determine the oligomerization state of the ATP synthase. In gel ATPase activity and western blotting for the ATP synthase β subunit in isolated mitochondria revealed a significant decrease in the dimer/monomer ratio and a total loss of oligomers in the *Tet3* knockout fibroblasts (Fig. 4d, e). Indeed, F_1_F_0_-ATP synthase showed a 60% reduced activity upon TET3 loss, as measured by spectrophotometry (Fig. 4f). Next, we evaluated whether an ATP synthase activity reduction might contribute to mitochondrial hyperpolarization. The inner mitochondrial membrane potential was significantly increased in the absence of TET3 (Fig. 4g). An increase in mitochondrial membrane potential upon ATP synthase assembly defect has been previously described^21^. The mitochondrial mass remained unaffected, as measured by confocal microscopy (Fig. 4g and Supp. Fig 3 b-e). The spectrophotometric quantification of the citrate synthase activity also did not show any differences between both genotypes (Fig. 4h). Finally, we found that the cellular respiration status (basal oxygen consumption rate or OCR) using an Agilent Extracellular Flux analyzer to measure basal oxygen consumption rate was not affected by the absence of TET3 (Fig. 4i). Overall, our data demonstrate that TET3 loss-of-function leads to a reduction in mitochondrial F_1_F_0_-ATP synthase activity and, consequently, to a hyperpolarization of the mitochondrial membrane potential that might be detrimental to the cell.

**Fig. 4.**
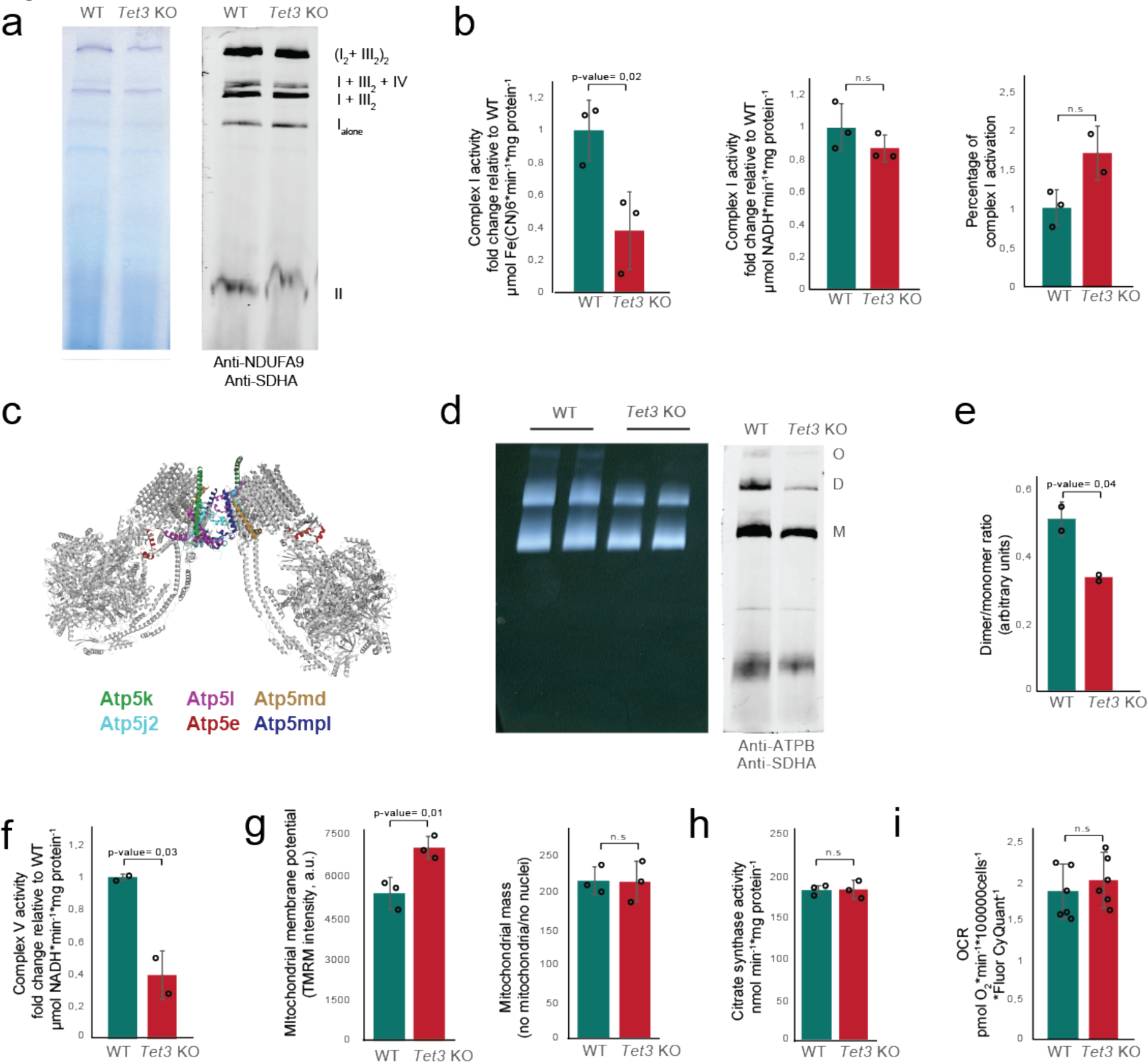
TET3 regulates mitochondrial F_1_F_0_-ATP synthase assembly. **a** Blue-Native gel electrophoresis followed by in gel complex I activity or western blot with anti-NDUFA9 and anti-SDHA in isolated mitochondria from wild-type and *Tet3* knockout fibroblasts. n=2 biologically independent replicates, each sample correspond to a pool of two independent biological samples. **b** Mitochondrial complex I NADH: Fe(CN)6 oxidoreductase (left panel) and NADH:ubiquinone (middle panel) activities spectrophotometrically measured in isolated mitochondria from wild-type and *Tet3* knockout fibroblasts. (right panel). Percentage of complex I activation in isolated mitochondria. n=3 biologically independent samples. **c** Bovine ATP synthase dimer state1:state3 model (PDB ID: 7AJD) in which the subunits showing lower expression levels in the absence of TET3 have been color-coded. **d** Clear native gel electrophoresis followed by in gel ATPase activity or blue native gel electrophoreses followed by western blot with anti-ATPβ from isolated mitochondria in wild-type and *Tet3* knockout fibroblasts. ATP synthase O, oligomer; D, dimer; M, monomer. n=2 biologically independent replicates, each sample correspond to a pool of two independent biological samples. **e** Dimer/monomer ratio quantification detected by in gel activity in (b). n=2 biologically independent replicates, each sample correspond to a pool of two independent biological samples. **f** Complex V activity measured spectrophotometrically in mitochondria isolated from wild-type and *Tet3* knockout fibroblasts. n=2 biological replicates. **g** Mitochondrial membrane potential and mitochondrial mass quantification in wild-type and *Tet3* knockout fibroblasts, as measured by confocal imaging. n=3 biologically independent samples. a.u., arbitrary units. **h** Citrate synthase activity measured spectrophotometrically in wild-type and *Tet3* knockout fibroblasts. n=3 biological replicates. **i** Oxygen consumption rate (OCR)-derived quantification of basal respiration in wild-type and *Tet3* knockout fibroblasts. n=6 biologically independent replicates. Error bars represent standard deviation. Student’s *t* test, two-tailed.

### Mitochondrial function is dependent on TET3 catalytic activity

We next search for aberrantly methylated genes with relevant mitochondrial functions by defining 5mC and 5hmC global patterns using reduced representation oxidative bisulfite sequencing (RRoxBS-seq). Our analysis revealed that the overall degree of DNA methylation levels in promoters, CpG islands as well as along the gene body regions and other regulatory regions was higher in the *Tet3* knockout in comparison with the wild-type (Fig. 5a, b). Plotting only the significant differentially methylated CpGs across the gene body highlighted a specific *Tet3* knockout signature with a 5m CpG hypermethylation peak around the upstream body region and a hypomethylation peak at the downstream body region (Fig. 5c). However, no differences were observed in 5hm CpG levels between the wild-type and the *Tet3* knockout (Fig 5c). As TET3 impairment leads to hypermethylation, we examined differentially methylated regions (DMRs) between the *Tet3* knockout and the wild-type. Our analysis showed that hypermethylated DMRs are dominant and located in introns and distal intergenic regions (Fig 5d, e). At a single-nucleotide level, DNA methylation was selectively increased in *Atp5e*, which negatively correlates with its transcription level as was shown in Fig. 2g. Indeed, *Atp5e*, a gene that encodes a protein located in the F_1_F_0_-ATP synthase dimerization domain and is required for its dimmerization, exhibits a DMR with a significant higher methylation level than the control (72% vs 36%, respectively) (Fig 5 f, g). Therefore, a compromised *Atp5e* gene expression due to increase methylation levels in the absence of TET3 promotes a substantial reduction in the formation of F_1_F_0_-ATP synthase dimmers and oligomers, resulting in mitochondrial alterations.

**Fig. 5.**
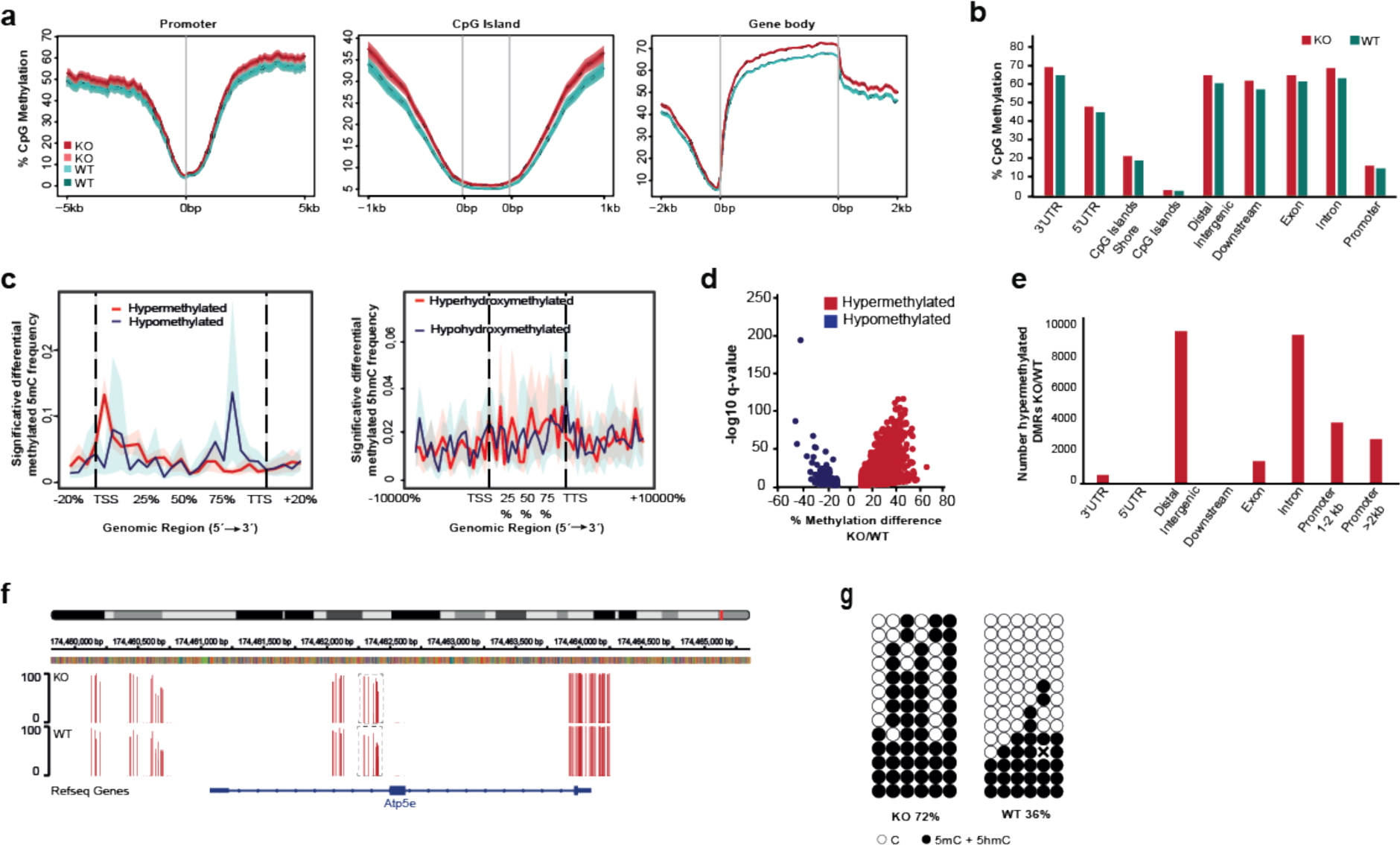
TET3 deletion leads to hypermethylation. **a** RRoxBS data represented as 5mC average levels (100 bp-bin) over promoters, CpG islands and gene bodies for all annotated genes in *Tet3* knockout and wild-type fibroblasts. n=2 biological replicates. Standard error and 95% confidence interval are represented as dark and light shaded areas around the mean line, respectively. **b** *Tet3* knockout and wild-type fibroblast methylation frequencies across different genomic elements. n=2 biological replicates. **c** Distribution of CpG sites with significantly different methylation (q-value <0,05) between *Tet3* knockout and wild-type, determined from RRoxBS. n=2 biological replicates. Standard error is denoted as a shaded area around the mean line. **d** Differential DMR methylation levels between *Tet3* knockout and wild-type. The blue and red dots represent hypo- and hypermethylated DMRs according to the definition described in methods and P<X n=2 biological replicates. **e** Distribution of hyper-DMRs present in *Tet3* knockout fibroblasts across various genomic features. n=2 biological replicates. **f** Gene tracks showing RRoxBS for *Tet3* knockout and wild-type cells over *Atp5e* genomic region. One of two representative biological replicates is shown. Vertical bars above the horizontal line indicate the methylation level (0-100) at individual CpG. The dash line box depicts a hyper-DMR in one Atp5e intron. **g** Bisulfite sequencing analysis of the *Atp5e* intron in *Tet3* knockout and wild-type fibroblasts. The percentages of methylated + hydroxymethylated CpGs are indicated.

### TET3 loss-of-function impairs mitochondrial metabolic maturation

To assess the impact of the ATP synthase assembly deficiency on cellular metabolism, we performed a metabolic analysis targeted on central carbon metabolism. First, we quantified the redox couples nicotinamide adenine dinucleotide phosphate (NADP(H)), nicotinamide adenine dinucleotide (NAD(H)) as well as glutathione (GSH) and oxidized glutathione (GSSG) by mass spectrometry (Fig. 6a). In the absence of TET3, we observed an increase in NADH as well as in NAD+ and its precursor NAAD (Fig. 6c). NADPH is required for fatty acid synthesis and for maintaining the steady-state level of GSH. NADPH is slightly but not significantly increased in the absence of TET3 whereas NADP+ is significantly elevated. GSH is one of the most important non-enzymatic antioxidants. There is an increment in the levels of both GSH and GSSG. FAD was also increased in the absence of TET3 (Fig. 6b). Next, we observed a significant increment in the levels of almost all metabolites from glycolysis (Fig. 6d) and a slight but significant rise in lactic acid was measured in the TET3—depleted fibroblasts (Fig. 6f). Furthermore, all the metabolites of the pentose phosphate pathway (PPP), which plays a pivotal role in the production of cellular NADPH and nucleotide biosynthesis, were elevated (Fig. 6d). In fact, both pyrimidine and purine nucleotides concentrations were increased in the absence of TET3 (Fig. 6e). The metabolites involved in hexosamine pathway and tricarboxylic acid (TCA) cycle were also higher upon loss of TET3 function (Fig. 6d). The levels of acetyl-CoA that is required for fatty acid synthesis were also augmented in the *Tet3* knockout fibroblasts (Fig. 6d). In addition, higher levels of several glycerophospholipids, which are the major structural lipids in eukaryotic membranes^22^, such as glycerophosphoetanolamine, glycerophosphoserine, glycerophosphoglycerol and glycerophosphoinositol, were also detected (Fig. 6g). Moreover, acetoacetyl CoA that is essential for cholesterol synthesis also exhibited a significant increase in TET3—deployed cells (Fig. 6h). GDP glucose/mannose, a sugar donor for carbohydrate biosynthesis, displayed an increment in the absence of TET3 as well (Fig. 6i). Some of the enzymes involved in these metabolic pathways show substrate promiscuity and produce side products that might be toxic ^23^. A significant 2-fold increase of 4-phosphoerythronate, the resulting side product generated by glyceraldehyde 3-phosphate dehydrogenase acting on erythrose-4-phosphate instead of on glyceradehyde-3-phosphate, was measured in *Tet3* knockout fibroblasts. Likewise, 2-hydroxyglutarate that arises from the noncanonical activity of several dehydrogenases also accumulates in TET3-depleted fibroblasts as well as methyl-malonic acid that is a byproduct of propionate metabolism. The accumulation of these side products correlates with the observed increase in the glycolytic and TCA flux (Fig 6j). Phosphocreatine, which is considered an energy buffer^24^ as well as ATP were increased in *Tet3* knockout fibroblasts (Fig. 6e, k). Overall, our analysis indicates that TET3-deployed cells are enriched in glycolysis-dependent anabolic pathways that are reminiscent of undifferentiated cells^25^. These results are consistent with the observed immature mitochondrial morphology and impaired oxidative phosphorylation, suggesting that the loss of TET3 impedes mitochondrial maturation and compromises terminal differentiation at the metabolic level.

**Fig. 6.**
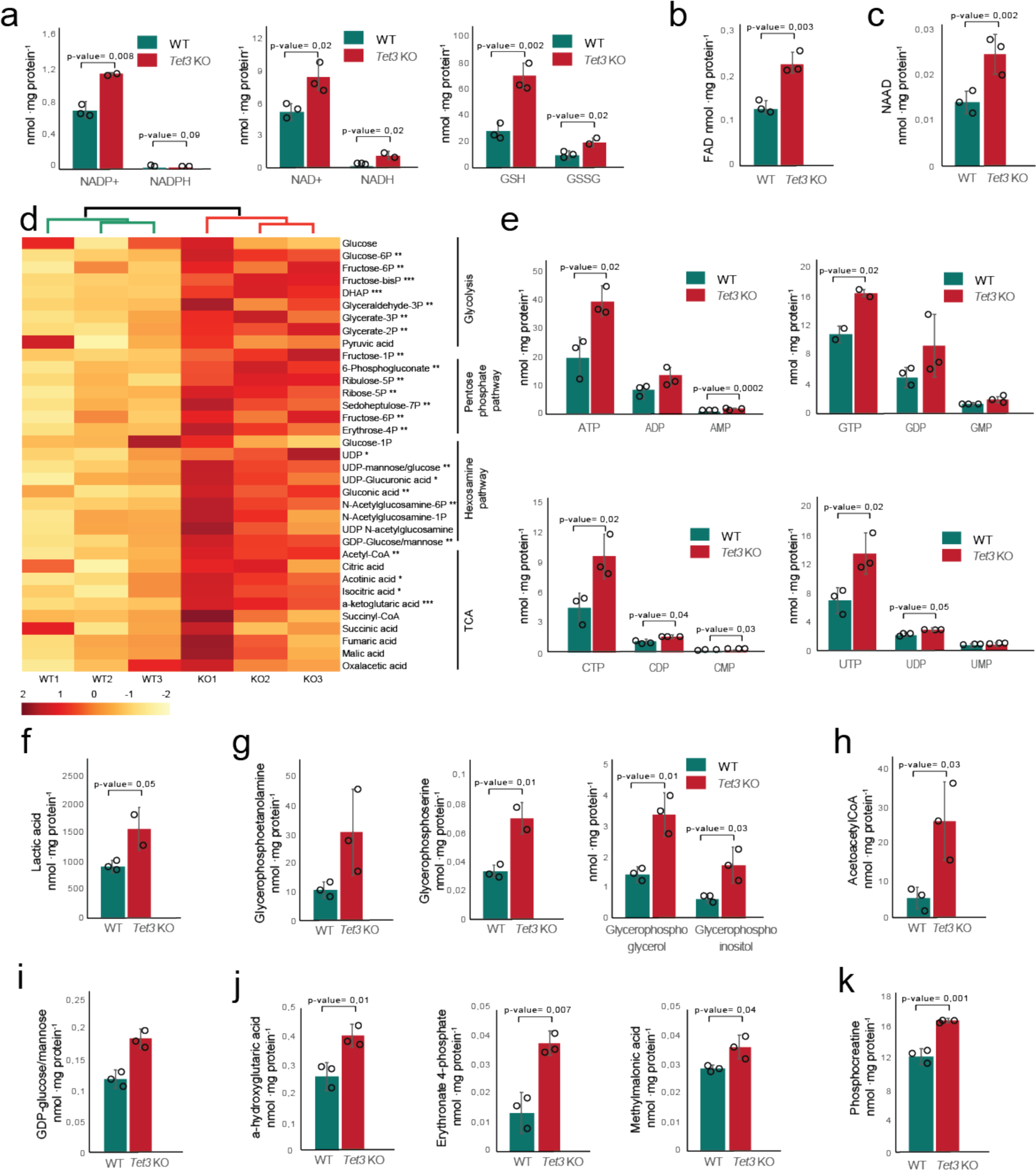
TET3 regulates terminal differentiation at the metabolic level. **a-c** Mass spectrometry quantification of the following metabolites: **a** NAD(P)H, NAD(P)+, GSH and GSSG, **b** FAD, **c** NAAD. **d** Heatmap visualization of the z-scored levels from glycolysis, pentose phosphate pathway, hexosamine pathway and tricarboxylic acid pathway metabolites in wild-type and *Tet3* knockout fibroblasts measured by mass spectrometry. Color represent row z-score of each sample. **e-k** Mass spectrometry quantification of the following metabolites: **e** nucleosides, **f** lactic acid, **g** glycerophospho-etanolamine, -serine, -glycerol and -inositol, **h** acetoacetylCoA, **i** GDP-glucose/mannose, **j** α-hydroxyglutaric acid, erythronate-4-phosphate, methylmalonic acid, **k** phosphocreatinine. n=3 independent biological replicates. Error bars correspond to the standard deviation. Student *t* test, two-tailed.

## DISCUSSION

The epithelium of the small intestine represents a valuable model to study stem cell regulation due to fast and continuous cycles of self-renewing, differentiation and apoptosis. A metabolic gradient can be observed along the crypt-villus axis in which the stem cells located in the crypt depend mainly on glycolysis for energy supply whereas enterocytes dependence on oxidative phosphorylation increases with differentiation and maturation while migrating upwards to the lumen^7^. In this study, we have demonstrated that, in the absence of TET3, enterocytes cannot acquire a physiological phenotype due to a stalled mitochondrial metabolic maturation that prevents terminal differentiation at the metabolic level.

Two recent reports using an intestinal epithelium—specific *Tet3* or a double *Tet2* and *Tet3* conditional knockout mouse model concluded that TET3 is essential for proper epithelial differentiation but the underlying molecular mechanism was not defined^26,27^. It has also been proposed that TET3 ablation predisposes the intestinal epithelium to inflammation through microbiome alterations^27^. Likewise, another study has predicted TET enzymes to be risk genes in ulcerative colitis patients^28^. Our results suggest that the production of reactive species due to an ATP synthase assembly defect might also contribute or even be the major cause leading to inflammation in these pathologies. Furthermore, a study in the cardiac tissue showed that a cardiac-specific deletion of both *Tet2* and *Tet3* in mice leads to ventricular non-compaction cardiomyopathy with embryonic lethality^13^. Our findings can explain this cardiac phenotype since patients exhibiting ATP synthase deficiency due to single mutations in other ATP synthase subunits or assembling factors such as *ATP5F1D* or *TMEM70* display cardiomyopathies^29,30^. Recently, a human mendelian disorder caused by the disruption of *TET3* has been reported^31^. However, an insight into the disease mechanism was not described. We propose that the pathology observed in TET3 deficient patients might be due to a compromised complex V dimmerization, as none of the described *TET3* mutations induce a complete TET3 loss-of-function^31^, and point to *Tet3* as a new candidate gene to be considered for F_1_F_0_-ATP synthase deficiency diagnosis.

TET catalytic loss-of-function strongly associates to tumorigenesis since most human cancers exhibit a dramatic loss of 5hmC when compared to their normal tissue counterparts^32^. Although TET mutations can be found in some tumor types such as myeloid malignancies, gliomas and a few others, most human cancers do not contain mutations on the TET genes or their co-factors, and yet, almost every tumor analyzed to date exhibits a clear loss of 5hmC^32^. However, it is important to keep in mind that TET enzymes can be enzymatically inhibited due, for instance, to hypoxia conditions or the presence of oncometabolites. Loss-of-function mutations in TET2 are associated with diverse myeloid and lymphoid leukemia in humans, however, these mutations *per se* are not sufficient to induce tumorigenesis and a second unknown hit is required. *Tet2* knockout mice develop lethal myeloid malignancies at around 1-year of age^9^. Interestingly, a conditional double *Tet2* and*Tet3* knockout mouse model showed a very aggressive leukemia between 3 and 7-weeks of age but the molecular mechanism underlying this oncogenic transformation was not defined^33^. An et al. suggested that TET2-depletion leads to an impairment in double stranded break repair mechanism ^33^. Our findings demonstrate that TET3 loss establishes a metabolic program in which oxidative capacity is reduced whereas glycolysis-dependent anabolic pathways are enhanced due to mitochondria with an immature phenotype. This metabolic signature resembles the metabolic profile observed in undifferentiated cells^25^ and suggests that TET3-depleted cells might be primed to malignancy transformation. Mitochondrial hyperpolarization due to compromised F_1_F_0_-ATP synthase dimerization and activity can generate an excess of reactive oxygen species and induce oxidative damage^21^. Hence, we hypothesize that a deficiency in double stranded break repair mechanisms combined with an increase in oxidative stress might lead to genomic instability and oncogenic transformation in cells that, in addition, display a metabolic profile closely related to undifferentiated cells. This scenario might explain the extremely fast development of leukemias observed in the conditional double *Tet2* and*Tet3* knockout mouse model. Collectively, our analysis has revealed the molecular mechanism by which TET3 function regulates terminal differentiation at the metabolic level and by which its loss-of-function might contribute to tumorigenesis initiation together with other factors such as TET2 or others still not defined. Future work will need to assess whether our findings can be extrapolated to all cell-types of the organism since *Tet3* is ubiquitinously expressed in differentiated cells.

## METHODS

### Generation of *Tet3* knockout mice

A targeting vector introducing *loxP* sites around *Tet3* exon 5 was constructed and electroporated into (C57BL6/J x C3H/HeJ) F1 embryonic stem cells (ESCs). Deletion of the floxed region results in a frame-shift that introduces a premature stop codon in exon 6, triggering nonsense-mediated decay of the mutant transcript. The construct also contained an *FRT*-flanked selection cassette coding for neomycin and thymidine kinase. Individual neomycin-resistant ESC clones were screened for recombination in the proper genomic locus. Next, FIAU-negative selection was used to select for clones that had excised the thymidine kinase cassette after transient transfection of an Flp-coding plasmid. Then, a Cre-coding plasmid was transiently transfected to excise the *loxP*-flanked exon 5 and single ESC clones were genotyped to confirm the excision. *Tet3* knockout ESCs were used to generate chimeric mice through aggregation with morula stage (C57BL6/J x C3H/HeJ) F1 embryos. The generated male chimeric mice were crossed to CD1 females to screen for germ line transmission of the *Tet3* knockout allele. Finally, Tet3 heterozygous females were backcrossed with pure C57BL6/JRccHsd males at least 5 generations to obtain an incipient congenic line (N5 or >95% C57BL6/JRccHsd). Genotyping primers are shown in Supp. Table 5. Mice were time-mated and the presence of a vaginal plug was considered as embryonic day (E) 0.5. Experiments were performed on E18.5 embryos obtained by caesarian section, as the gestation length of this particular mouse line is 19 hours. Animals were housed in a controlled environment with a 12h light/dark cycle in open cages, with access *ad libitum* to a standard chow diet and water. All animal experiments were performed in accordance with the animal care guidelines of the European Union (2010/63/EU) prior approval of the Spanish National Research Council Ethics Committee and the Regional Government competent authority.

### Immunofluorescence, confocal microscopy and image analysis

Small intestine was dissected from E18.5 embryos, fixed by immersion in 4% paraformaldehyde, dehydrated through increasing ethanol concentrations, cleared in xylene and embedded in paraffin blocks from which 5 µm sections were obtained. Double stainings were performed first for 5hmC detection using the Tyramide Superboost kit (ThermoFisher Scientific) followed by stripping and second staining for cell-type specific markers or 5-methylcytosine. Briefly, sections were deparaffinized, rehydrated and permeabilized for 30min in 0.5% Triton X-100. Antigen retrieval was performed by incubating sections 25 min in sodium citrate buffer 10mM pH 6 in a water bath at 95°C. Next, 5hmC staining was performed following the Tyramide Superboost kit recommendations. Sections were subsequently stripped using sodium citrate buffer as described above and blocked for 1 h in blocking solution consisting of 2% normal goat serum, 2% bovine serum albumin (BSA) and 0.2% Triton X-100 in PBS. Sections were then incubated overnight at 4°C with primary antibody, 1 h at room temperature with secondary antibody and finally 5 min with DAPI. Antibodies are listed in Supp. Table 6. Images were acquired with a SP8 confocal microscope (Leica Microsystems) and processed using the Fiji software. For cell counting, three pairs of E18.5 wild-type and *Tet3* knockout small intestines, each pair from the same litter, were used. Each sample was stained for 5hmC together with each cell-type specific marker and 5hmC—positive cells were counted using the Fiji software^34^. The number of 5hmC—positive cells was calculated as a percentage of the total number of positive cells for each cell-type specific marker. Whole embryo images were obtained with an Aperio VERSA Scanner (Leica Microsystems).

### Intestine dissociation and cell sorting

A mouse small intestinal epithelium single cell suspension was performed based on a previous reported protocol^35^. Briefly, the small intestine was dissected from E18.5 embryos, opened longitudinally, and placed on ice-cold PBS containing 30mM EDTA and 1.5mM DTT for 20min on ice. Next, the intestine was transferred to a tube containing 30mM EDTA in PBS, incubated at 37°C for 10min and, then, vigorously shaken for 30s to release the intestinal epithelium from the lamina propia. After removing the intact intestinal muscle, the villi suspension was used for 5hmC genome wide profiling and targeted metabolite analysis or dissociated into single cell for fluorescence activated cell sorting. To generate a single cell suspension, the villi were incubated at 37°C for 14min in 10ml HBSS (with Ca^2+^ and Mg^2+^) containing 8mg of dispase II. During the incubation, the tube was vigorously shaken for 30s every 2 min to dissociate the villi. Finally, 10% fetal bovine serum (FBS) and 0.5mg DNase I were added to the cellular suspension that was sequentially passed through 100 µm and 40µm filters for obtaining single cells. The single cell suspension was incubated in PBS containing 3% FBS and the CD31-PE, CD45-PE, Ter119-PE, EpCAM-APC780 fluorescent-labelled antibodies for 30min. DAPI was used for staining dead cells. A BD FACS Aria Fusion (BD Biosciences) was used for sorting the live intestinal epithelial population (DAPI—negative, CD31-PE—negative, CD45-PE—negative, Ter119-PE—negative and EpCAM-APC780—positive cells). For all fluorescent channels, positive and negative cells were gated on the basis of fluorescent minus one control. Antibodies are listed in Supp. Table 6. For RNA applications (quantitative RT-PCR and scRNA-seq), all steps were performed under RNase-free conditions and 0.5U/µl of RiboLock RNase inhibitor (ThermoFisher Scientific) was added to all buffers.

### 5hmC and 5mC quantification

DNA was extracted from sorted intestinal epithelial cells using the Qiamp DNA Mini kit following the manufacturer’s protocol. DNA was hydrolysed for 6 h at 37°C using DNA degradase plus (Zymo Research). Mass spectrometry quantification was performed as previously described^36^ in the Babraham Institute mass spectrometry facility using three individual E18.5 wild-type and three individual E18.5 *Tet3* knockout embryos.

### RT - quantitative PCR

Sorted intestinal epithelial cells from three different wild-type or *Tet3* knockout embryos were combined and RNA was extracted using the Qiamp RNeasy micro kit, including the DNAse treatment step, following the manufacturer’s protocol. cDNA was obtained by reverse transcription using the M-MLV Reverse transcriptase and quantitative PCR was performed in technical triplicates on a Quant Studio 5 Real-Time PCR System (Applied Biosystems) using iTaq Univeral SYBR Green Supermix (Biorad) and a primer concentration of 375nM. Primers are listed in Supp. Table 5. *Actb* and *Gapdh* were used as endogenous controls for gene expression analysis. Fold enrichment normalized to wild-type samples was calculated using the 2^-ΔΔCt^ method^37^.

### Single cell RNA-seq

Sorted E18.5 intestinal epithelial cells were fixed in 90% ice-cold methanol and stored at −80°C until analysis. *Tet3* knockout and wild-type samples correspond to a pool of intestinal epithelial cells from two embryos each. 6,000 cells per sample were analyzed. For analysis, cells were resuspended in 3x saline sodium citrate buffer containing 0.04% BSA, 1% SUPERase·In RNAs Inhibitor (Invitrogen) and 40 mM DTT^38^. Single-indexed libraries were prepared using the Chromium Single Cell 3’ v3/v3.1 chemistry (10x Genomics) following the manufacturer’s instructions. The generated libraries were sequenced on an Illumina Novaseq (Illumina) system using the following transcript read lengths: 28bp for cell barcode and UMI, 8bp for sample index and 91bp for insert. 10,000 reads per cell were sequenced. scRNA-seq analysis was performed at Princess Margaret Genomics Centre (Toronto, Canada).

### Single cell RNA-seq data analysis

For scRNA-seq data processing, the alevin pipeline integrated with the salmon software (version salmon-0.14.1) was used, performing cell barcode (CB) detection, read mapping, unique molecular identifier (UMI) deduplication, gene count estimation and cell barcode whitelisting^39^. Reads were aligned and annotated to the mouse GRCm38 reference genome, transcriptome and annotation, downloaded from GENOME website (version 21). Alevin’s forceCells option was set to 6,000 for both wild-type and *Tet3* knockout populations to specify the number of CBs to consider for whitelisting. Standard procedures for filtering, variable gene selection, dimension reduction and clustering were performed using the Scanpy toolkit (version 1.7.1)^40^. Cells with fewer than 200 detected genes were removed from the analysis as well as cells with a mitochondrial versus all gene fraction greater than 0.15. Genes detected in more than 3 cells were considered as expressed, while genes with more than 100,000 total counts were excluded. The data matrix was total-count normalized to 10,000 reads per cell and stabilized by computing the natural logarithm of one plus the counts for each cell. Highly variable genes were filtered for feature selection, remaining a data matrix of 11,871 cells and 11,871 genes. Each gene was scaled to unit variance and values exceeding standard deviation 10 were clipped. For dimensional reduction and clustering, the t-Distributed Stochastic Neighbor Embedding (t-SNE)^41^ and the Louvain graph-clustering method^21^ were used. The ranking for the characterizing genes of each cluster was computed using the Wilcoxon test with Benjamini-Hochberg correction method. For trajectory inference, a partition-based graph abstraction (PAGA) map preserving the global topology of data was performed^40^ and single-cell graphs using ‘fa’ (ForceAtlas2)^42^ layout and PAGA initialization were made, one for the Louvain cluster groups, and another for the wild-type and *Tet3* knockout groups. Gene ontology analysis was performed using the *g:Profiler* web server with default parameters^43^ using the top 100 gene markers for wild-type or *Tet3* knockout specific clusters. Gene Set Enrichment Analysis was performed with GSEA v.4.1.0 desktop software using the GSEA Preranked tool with SIGN(log2FC)*(-log10(p-value)) as ranking parameter for all detected genes and c2.cp.kegg.v7.4.symbols.gmt as enrichment gene set^44^.

### Transmission Electron Microscopy

Brain, heart and intestine were obtained from *Tet3* knockout and wild-type 18.5 d.p.c. embryos obtained by caesarian section. Tissues were immediately fixed overnight at 4 °C by immersion in 0.1 M phosphate buffer pH 7.4, containing 4% paraformaldehyde and 2% glutaraldehyde, followed by postfixation with 1% osmium tetroxide-0,8% potassium ferrocyanide mixture in water for 1 hour at 4 °C. Samples were then dehydrated with ethanol and propylene oxide and embedded in Epon resin. Semi-thin sections were obtained from the resin blocks and stained with toluidine blue to identify regions of interest in intestine and brain. Ultra-thin sections were stained with lead citrate and examined in a HT780 Hitachi electron microscope. Sample preparation and imaging were generated in the Electron Microscopy Service (University of Valencia). Images were analyzed with ImageJ v.1.6.0.

### Mitochondria isolation

Isolation of mitochondria from cell cultures was performed from 4-10 150 mm plates, according to the differential centrifugation method. Briefly, cells were lysed by hypotonic shock with 7 volumes of hypotonic medium (83mM sucrose, 10mM MOPS, pH 7.2) and homogenized in a Teflon potter-type tissue homogenizer. Then, 7 volumes of hypertonic medium (sucrose 510, MOPS 30mM, pH 7.2) were added. The mixture was centrifuged at 1000g, 5 minutes and the supernatant obtained at 12000 g, 12 minutes. The pellet obtained was resuspended in medium A (0.32M sucrose, 1mM EDTA, 10mM Tris-HCl, pH 7.4). After several centrifugations at 15000g, 2 minutes, until a clear supernatant was obtained, the pellet was resuspended in medium A (0.32M sucrose, 1mM EDTA, 10mM Tris-HCl, pH 7.4). The pellet was resuspended in medium A and stored at −80°C.

### Mitochondria solubilization

Solubilization of mitochondrial membranes was carried out with digitonin (DIG), in order to visualize both respiratory SCs and free form complexes. The ratio of grams of detergent: grams of mitochondrial protein used was 4:1, except in gel where complex V was visualized in which the ratio was lowered to 2.5:1. To 100 μg of purified mitochondria, resuspended in 50mM NaCl buffer, 50mM imidazole, 5mM aminocaproic acid at a concentration of 10 μg/μl, digitonin was added at the desired concentration. Mitochondria were resuspended, at this point the solution characteristically become clearer for the effect of the detergent, and incubated 10 minutes on ice. The insoluble portion was removed by centrifugation at 13000 g in microfuge for 30 minutes at 4oC. The pellet obtained was discarded and the supernatant was mixed with mixed with 4x loading buffer (5% Coomassie Blue-G250 in 1M aminocaproic acid) for gel loading.

### Blue Native-PAGE and Clear Native-PAGE

3-13% gradient polyacrylamide gels were prepared in house with a gradient former. CN gels were essentialy the same as BN gels, but 0.01% of digitonin was added to all gel solutions and Ponceau red buffer (Ponceau red, glycerol) was used instead of the normal loading buffer. The amount of sample loaded in each well was that obtained from solubilization of 100 μg of mitochondria. Cathode buffer A (tricine 50mM, bis-tris 15mM, pH 7.0, Coomassie Blue G-250 0.02%), cathode buffer B (tricine 50mM, bis-tris 15mM, pH 7.0, Coomassie Blue G-250 0.002%) and anode buffer (bis-tris 50mM, pH 7.0) were used for electrophoresis. Electrophoresis was performed in cold chamber. The first half hour was run at 90 volts with cathode buffer A. After that time, the cathode buffer was exchanged for cathode buffer B. Electrophoresis continued for approximately one more hour at 300 volts, until the dye began to run out of the bottom of the gel. In CN gels the whole electrophoresis was performed in cathode buffer B, with no changes.

### In-gel complex I activity

Measurement of NADH dehydrogenase activity of complex I was determined on the same gel after BN-PAGE electrophoresis. The gel was incubated in 0.1 M Tris-HCl, pH 7.4, 0.14 mM NADH and 1 mg/ml NitroBlue tetrazolium solution at room temperature. Visualization was achieved by the purple precipitate produced after reduction of NitroBlue tetrazolium by the NADH dehydrogenase activity of CI.

### In-gel complex V activity

Measurement of NADH dehydrogenase activity of complex V was determined on the same gel after CN-PAGE electrophoresis85. The gel was incubated in 270 mM glycine, 35 mMTris, pH 8.4 for 3 hours, then ATP hydrolysis was assayed in 270 mM glycine, 35 mMTris, 4 mM ATP, 14 mMMgSO4, 0.2%Pb(NO3)2, pH 8.4. The reaction was stopped after 5–10 min by 30 min incubation in 50% methanol, 50% water after the formation of phosphate lead precipitates. The gel was then scanned.

### Spectrophotometric activities

Measurement of complex I activity involved a two-step measurement. Briefly, as described in^45^. First step was performed in 25 μg of 3x frozen-thawed mitochondria solubilized in Mg^2+^-containing C1/C2 buffer, containing DecylCoQ 130 μM and Antimycin A 1 μM. Absorbance measurement at 37°C, 340 nm started after addition of NADH 100 μM and lasted 2-4 min. After, rotenone 1 μM was added and absorbance was again measured for 2-4 min. 1 mM Fe(CN)_6_ was added and measurement was performed at 420 nm at 37° for 2-4 min. Later, Diphenyleneiodonium chloride (DPI) was added and absorbance was again captured in the same conditions.

Complex V activity was measured as a readout of its ATPase activity. As described in^46^, oligomycin-sensitive ATPase activity (i.e. complex V activity) was calculated after measuring changes in absorbance at 340 nm driven by the pyruvate kinase reaction coupled to ADP phosphorylation by complex V.

### Analysis of mitochondrial membrane potential

For imaging of mitochondrial membrane potential, wild-type MEFs and KO in TET3 were seeded onto 96-well plate (PerkinElmer) and mitochondrial inhibitors were added at Krebs Ringer Phosphate Glucose buffer (KRPG) supplemented with NaCl 145 mM; Na_2_HPO_4_ 5,7 mM; KCl 4,86 mM; CaCl_2_ 0,54 mM; MgSO_4_ 1,22 mM; glucose 20 mM; pH 7,35. Furthermore, cells were loaded with 10 nM TMRM (Sigma-Aldrich) and 1 µM cyclosporine-H (Sigma-Aldrich) in the Operetta CLS microscope (30 min at 37 °C in a 5 % CO_2_ atmosphere) and confocal images were acquired a 40X, 1.4 NA objective (PerkinElmer). Mitochondrial uncoupler carbonyl cyanide *p*-tri-fluoromethoxyphenylhydrazone (FCCP, 10 µM, 15 min, Sigma-Aldrich) was added as a control of mitochondrial depolarization. Finally, images were analyzed using Harmony software (PerkinElmer).

### Citrate synthase activities

E18.5 tail tip—derived fibroblasts from wild-type and *Tet3* knockout embryos were snap-frozen. After three cycles of freeze/thawing, to ensure cellular disruption, citrate synthase activities were determined. Citrate synthase activity was measured in the presence of 93 mM Tris-HCl, 0.1% (vol/vol) Triton X-100, 0.2 mM acetyl-CoA, 0.2 mM DTNB; the reaction was started with 0.2 mM oxaloacetate, and the absorbance was recorded at 412 nm (30 °C) (*e* = 13.6 mM^−1^ ·cm^−1^)^47^. The protein concentration of the samples was quantified by the BCA protein assay kit (Thermofisher) following the manufacturer’s instructions, using BSA as a standard.

### Oxygen consumption measurement

Oxygen consumption assays were performed as in PMID: 32728214. Briefly, oxygen consumption rate (OCR) was measured using an XF96 Extracellular Flux Analyzer (Seahorse Bioscience). MEFs were plated 1 day before the experiment and preincubated with unbuffered DMEM 1 h at 37 °C in an incubator without CO_2_ regulation. Successive injections of unbuffered DMEM, 5 μg/mL oligomycin, 300 nM carbonyl cyanide 4-(trifluoromethoxy) phenylhydrazone (FCCP) and 1 μM rotenone plus 1 μM antimycin A were programmed. Calculations were performed following the manufacturer’s instructions.

### Reduced Representation Oxidative Bisulfite Sequencing (RRoxBS-seq)

*Tet3* knockout and wild-type samples correspond to fibroblasts derived from the tail tip of 18.5 d.p.c. embryos as primary culture. Two lines of each genotyped were analyzed. Briefly, 5mC and 5hmC modifications were determined using oxidative bisulfite sequencing (oxBS-Seq) ^48^ in combination with reduced representation bisulfite sequencing (RRBS) ^49^. Briefly, ∼300 ngs of genomic DNA were digested with 100U of MspI (New England Biolabs), and cleaned up using QIAquick PCR purification columns (Qiagen). The digested DNA was spiked with 0.01% control DNA duplexes, provided by Cambridge Epigenetix (Cambridge, UK), containing C, 5-mC and 5-hmC bases at known positions. The control sequences are later analyzed after sequencing to give a quantitative assessment of the efficiency of oxidation and bisulfite conversion. Uniquely indexed libraries were generated using the Ovation Ultralow Methyl-Seq DR Multiplex with TrueMethyl oxBS kit and workflow (Tecan). Each library was split into two aliquots for chemistry as follows: 2/3 were oxidized and bisulfite converted (oxBS), the other 1/3 was subjected to a mock oxidation before bisulfite conversion. 2ul of the resulting material was assessed by qPCR to determine the optimal number of PCR amplification cycles. After amplification the libraries were normalized to 2nM and pooled according to the desired plexity, clustered at 6.5pM on single read flow cell and sequenced for 100 cycles on an Illumina NovaSeq. Illumina’s CASAVA 1.8.2 software was used to perform image capture, base calling and demultiplexing of raw reads to produce FASTQ files. Quality of the raw sequence reads was assessed with fastp (https://github.com/OpenGene/fastp). Raw reads were then adapter trimmed, aligned, and post-processed to produce methylation calls at base pair resolution using a previously described in-house pipeline using CUTADAPT instead of FLEXBAR to trim the reads ^50^. CpG sites from the post-processing results were interrogated for true methylation (5mC) and hydroxy methylation (5hmC), by subtracting the oxBS methylation call from the BS methylation call on the same CpG, and for differential methylation (q value < 0.05 and methylation percentage difference of at least 25%) using methylKit ^51^. 5mC DMR is defined as a region with at least 3 CpGs, at least 1 is significantly differentially methylated CpG (DMC with 15% methylation difference and a q value < 0.05) and the DMR has average differential methylation of at least 10% across all CpGs in the region. 5hmC DMR is defined as a region with at least 2 CpGs, at least 1 is significantly differentially methylated CpG (DMC with 9% hydroxymethylation difference and a q value < 0.05) and the DMR has average differential hydroxymethylation of at least 8 % across all CpGs in the region. Library preparation for RRoxBS libraries, sequencing, post-processing of the raw data, data alignment and methylation calls were generated at the Epigenomics Core at Weill Cornell Medicine (New York, USA). The average plots were generated using the Seqplots Bioconductor R package ^52^. The 5mC and 5hmC single peaks were displayed using the Integrative Genomics Viewer ^53^.

### DNA methylation analysis

Genomic DNA was extracted from tail tip-derived fibroblasts with a QIAamp DNA kit (Qiagen), treated with bisulfite using the EZ DNA Methylation-Direct kit (Zymo Research) and amplified with NZY Taq II 2x Green Master Mix (NZY Tech). PCR products of bisulfite-treated DNA were purified with a Gel Extraction kit (NZY Tech) and cloned into TOPO cloning kit (Thermofisher). Individual clones were sequenced by standard Sanger sequencing and data were analysed by the online tool QUMA ^54^.

### Metabolomic analysis

Cells were fixed with 80% methanol, at 100 μL per 1 million cells. Cells were lysed with the aid of two metal balls at 30 Hz on a MM 400 mill mixer for 3 min, followed by centrifugal clarification at 21,000 g for 10 min. The clear supernatants were collected for the following assays. The precipitated pellets were used for protein assay using a standardized BCA procedure. For NAD and nucleotide quantification, an internal standard (IS) solution containing 13C or H2-labeled NAD, NADH, AMP and ATP was prepared in 50% methanol. Serially diluted standard solutions of the targeted metabolites were prepared in 80% methanol. 10 μL of the clear supernatant of each sample or each standard solution was mixed with 40 μL of IS solution. 10 μL aliquots of the resultant solutions were injected into a C18 column to run UPLC-MRM/MS with (-) ion detection on a Waters Acquity UPLC system coupled to a Sciex QTRAP 6500 Plus MS instrument, using a tributylamine buffer solution and acetonitrile as the mobile phase for binary-solvent gradient elution. For GSH and GSSG quantification, an internal standard (IS) solution containing isotope-labeled GSH and GSSG was prepared in water. Serially diluted standard solutions of GSH and GSSG were prepared in 80% methanol. 20 μL of the clear supernatant of each sample or each standard solution was mixed with 80 μL of the IS solution. 10 μL aliquots of the resultant solutions were injected into a HILIC column to run UPLC-MRM/MS with (+) ion detection on an Agilent 1290 UHPLC system coupled to an Agilent 6495B QQQ MS instrument with the use of 0.1% formic acid in water and in acetonitrile as the mobile phase for binary-solvent gradient elution. For analysis of glucose, glucose-6P and mannose-6P, 50 μL of the supernatant of each sample was mixed with 50 μL of a solution of 13C6-glucose as internal standard, 100 μL of 25 mM AEC solution and 20 μL of acetic acid. The mixture was allowed to react at 60 oC for 70 min. After reaction, 300 µL of water and 300 µL of dichloroform was added. The mixture was vortex mixed and then centrifuged. The supernant was injected in 10-μL aliquots to run UPLC-MRM/MS with positive-ion detection on an Agilent 1290 UHPLC system coupled to a 4000 QTRAP mass spectrometer, as previously described ^55^. The analysis of the TCA cycle carboxylic acids was carried out as previously described ^56^. Briefly, 20 μL of the supernatant was mixed with 20 μL of a D-or 13-labeled analogue for each analyte as the internal standard, 20 μL of 200 mM 3-NPH solution and 20 μL of 150 EDC-6% pyridine solution. The mixture was allowed to react at 30 oC for 40 min. After reaction, the solution was diluted 3 fold with water. 20 μL was injected to run UPLC-MRM/MS with negative-ion detection on an Agilent 1290 UHPLC system coupled to a 4000 QTRAP mass spectrometer. Finally, quantification of other metabolites were performed as follows. 50 µL of each supernatant was mixed 50 µL of water containing 50 pmol/mL GTP-13C10 and 50 µL of chloroform. The mixture was vortex mixed for 1 min, followed by centrifugation. 50 µL of each clear supernatant was taken out and mixed with an equal volume of water. 10 µL was injected to run UPLC-MRM/MS with negative-ion detection using a custom-developed reversed-phase LC-MRM/MS method for binary-solvent gradient elution. Concentrations of the detected analytes in the above assays were calculated with internal standard calibration by interpolating the constructed linear-regression curves of individual compounds. The metabolomic analysis was performed in UVic Genome BC Proteomics Centre (Canada).

### Statistical analysis

Data, unless otherwise stated, is represented as mean ± standard deviation. Two-tailed t-test was used to assess significant differences between groups. A minimum of three biological replicates constitute each group.

## ACKNOWLEDGEMENTS

We are grateful for Dr. David Oxley and the Babraham Institute mass spectrometry facility for the cytosine modification analysis. N.T is funded by Agencia Estatal de Investigación (RYC-2014-16359; SAF2015-66549-R; PID2019-105920RB-100), Ayudas Fundación BBVA 2016 and Generalitat Valenciana (ACIF-2021-195). J.P.B. is funded by the NextGenerationEU/PRTR and Agencia Estatal de Investigación (10.13039/501100011033; PID2019-105699RB-I00; PID2022-138813OB-I00 and PDC2021-121013-I00); European Comission action HORIZON-TMA-MSCA-DN (ETERNITY,

101072759), and La Caixa Research Health grant LCF/PR/HR23/52430016.

## AUTHOR CONTRIBUTION

I.M. performed the immunostaining experiments; M.C.-C. and N.T. carried out the methylation analysis; C. G-C, M.C.-C- and N.T. performed the electron microscopy experiments; G.W. and N.T. generated the knockout mouse model; D.G. and M.J.A.-B. analyzed the single-cell RNA-seq data; C. G-C, A.C. and P.H.-A. performed and interpreted the mitochondrial activity and assembly experiments under the supervision of J.A.E.; D. J.-B. and I.M.-R. analyzed and interpreted the mitochondrial membrane potential under the supervision of J.P.-B.; N.T. designed the project, performed and interpreted most of the experiments, provided financial support and wrote the manuscript.

