## Supplementary Info for "TET3 regulates cellular terminal differentiation at the metabolic level"

**Supplementary Fig 1**

a

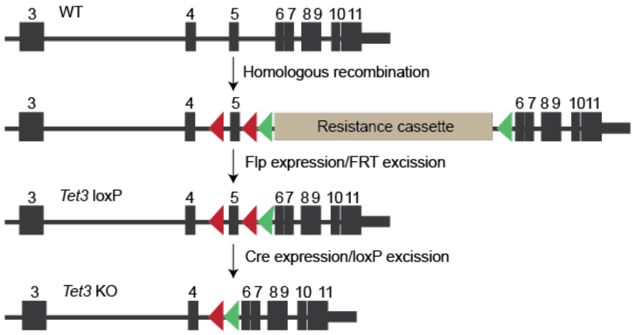


b

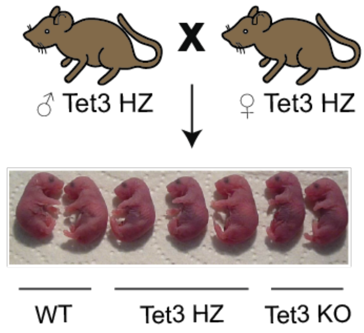


c

| | Mixed background | | Pure background | | Total | |
|---|---|---|---|---|---|---|
| Genotype | N embryos | % | N embryos | % | N embryos | % |
| WT | 66 | 29,9 | 55 | 22,4 | 121 | 25,9 |
| *Tet3* HZ | 103 | 46,6 | 134 | 54,5 | 237 | 50,7 |
| *Tet3* KO | 52 | 23,5 | 57 | 23,1 | 109 | 23,3 |
| Total | 221 | | 246 | | 467 | |

d

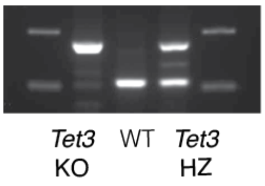


**Supplementary Fig. 1. a** Schematic representation of *Tet3* genomic locus and of the targeting strategy used to excise exon 5 in ESCs. Red and green head arrows represent *loxP* and *FRT* sites, respectively. **b** Representative litter of *Tet3* heterozygous mice showing no gross abnormalities in *Tet3* knockout newborns. **c** Summary of newborn genotypes showing normal Mendelian ratios for the *Tet3* knockout newborns. **d** Tail tip-PCR genotyping of wild type, *Tet3* heterozygous and *Tet3* knockout pups. WT, wild type; HZ, heterozygous; KO, *Tet3* knockout.

**Supplementary Fig 2**

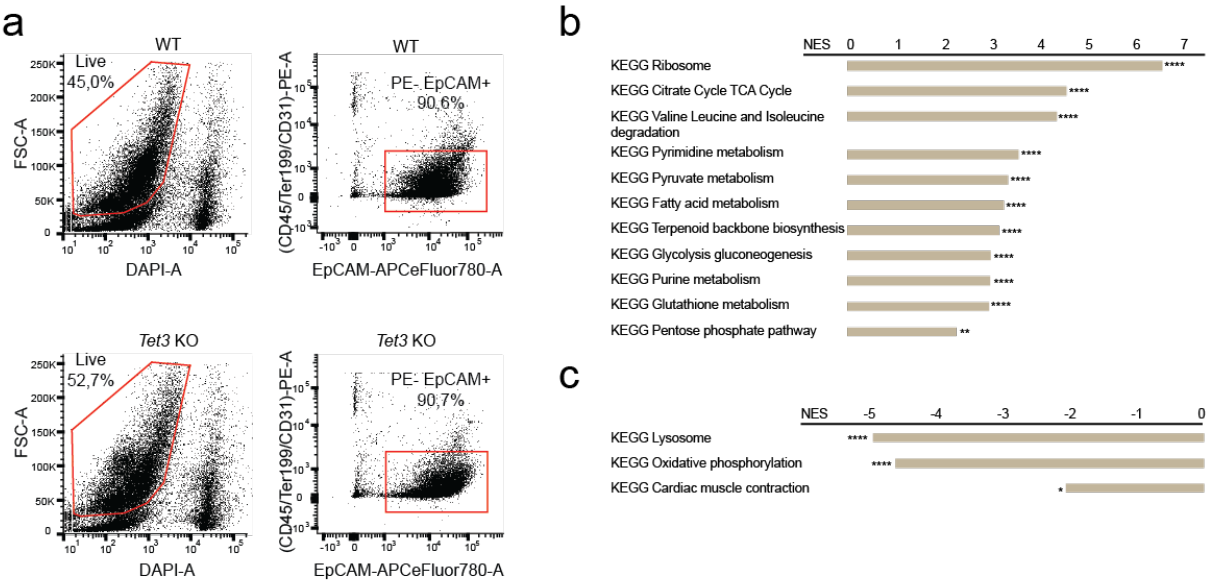


**Supplementary Fig. 2. a** Cell sorting strategy to sort live CD45PE—negative, Ter119PE—negative, CD31PE—negative, EpCAMAPCeFluor780—positive intestinal epithelial cells. (**b**, **c**) Top significantly positively **b** and negatively **c** enriched ontology terms identified using pre-ranked GSEA based on differences in gene expression between wild-type and *Tet3* KO specific clusters. NES, normalized enrichment score. ****FDR q-value = 0, **FDR q-value = 0,002, *FDR q-value =0,01.

### Supplementary Fig 3

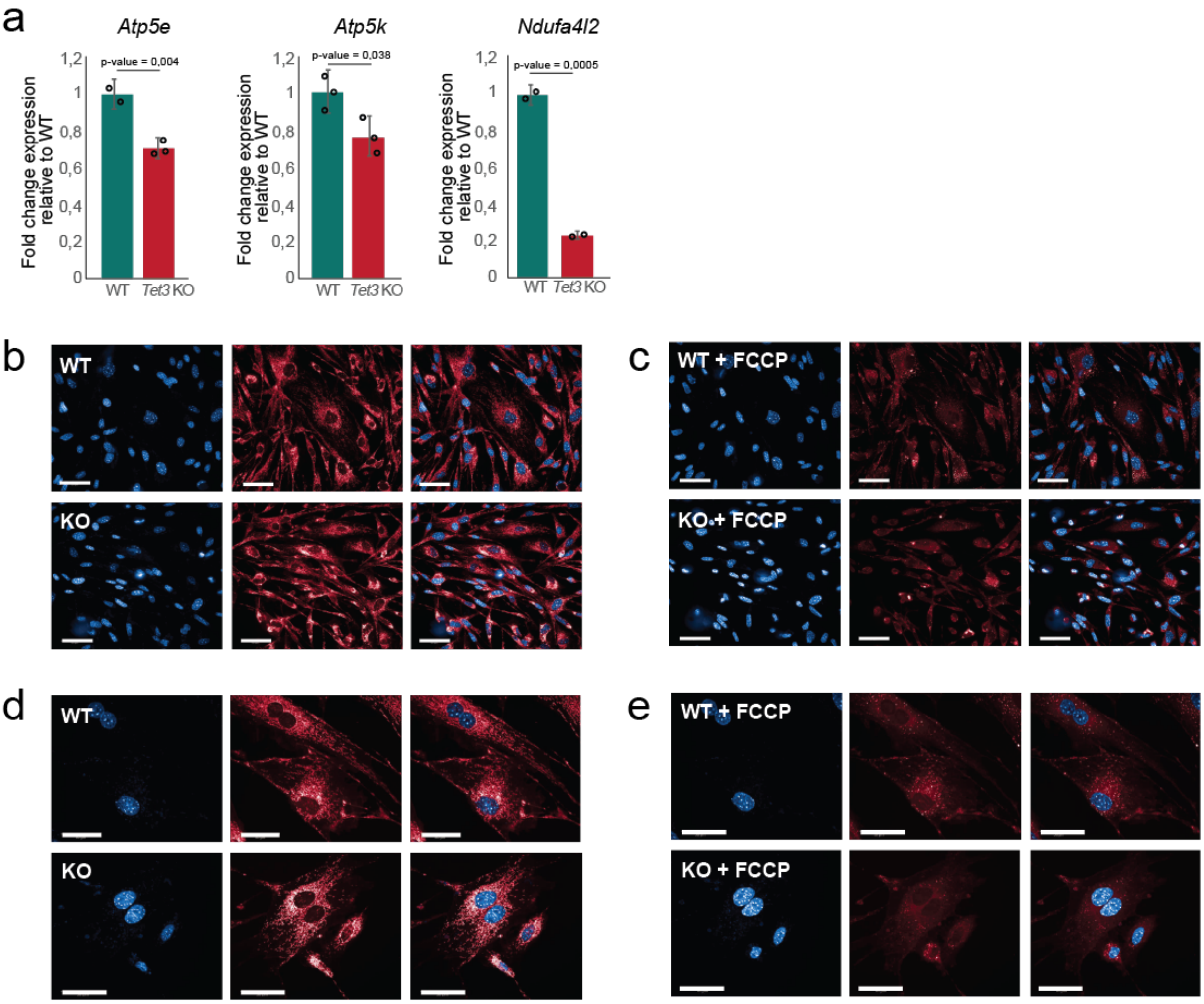


**Supplementary Fig. 3 a** Expression levels of one complex IV (*Ndufa4l2*) and two complex V (*Atp5e* and *Atp5k*) subunits as measured by RT-qPCR. Data has been normalized to the endogenous control and is shown as fold-expression relative to the wild-type that is considered as 1. Error bars represent standard deviation (n=3 technical replicates). **b**, **c** Representative confocal microscopy images after staining with TMRM (red) in wild-type and *Tet3* KO fibroblasts in the absence (**b**) or presence (**c**) of FCCP. The intensity of TMRM reflects the level of $\Delta\Psi m$. Hoechst was used to stain the nuclei (blue). (**d**, **e**) Higher magnification of panels **d** and **e**, respectively. Scale bar, 50 µm. Student's *t* test, two-sided.

**Supplementary Table 1** Small intestine epithelial gene markers used to define scRNA-seq cluster identity according to Haber *et al.* [1]

| Cell type | Markers |
|---|---|
| Stem/Transit-amplifying cells | *Acot1, C1qpbp, Ccdc34, Eef1a, H2afv, Hspd1, Naca, Nme1, Npm1, Ptma, Rpl13, Rpl22, Rpl32, Rpl37, Rps9, Rps15, Rps18, Rps19, Rps20, Siva1, Stmn1, Tomm5, Tubb5* |
| Enterocytes (proximal) | *Ace, Acsl5, Aldh1a1, Aldob, Apoa4, Apoc2, Apoc3, Cbr1, Cgref1, Ckb, Ckmt1, Cyb5r3, Dhrs1, Ephx2, Fabp1, Gda, Gpd1, Gpx4, Gstm1, Gstm3, H2-Q2, Lct, Leap2, Lpgat1, Mme, Ms4a10, Mttp, Prap1, Rbp2, Scp2, Slc2a2, Sult1b1* |
| Enterocytes (distal) | *Amn, Cndp2, Crip1, Cubn, Fabp6, Fgf15, Mep1a, Slc51a, Slc51b, Neu1,* |
| Goblet cells | *Agr2, Ccl6, Clca1, Fcgbp, Klk1, Muc2, Spink4, Tff3, Tpsg1, Zg16* |

**Supplementary Table 3** Wild type and *Tet3* knockout cell distribution in each cluster

| | **Wild Type** | **%** | ***Tet3* knockout** | **%** |
|---|---|---|---|---|
| **Cluster 0** | 1313 | 90 | 146 | 10 |
| **Cluster 1** | 1140 | 91,9 | 101 | 8,1 |
| **Cluster 2** | 7 | 0,6 | 1153 | 99,4 |
| **Cluster 3** | 10 | 1 | 968 | 99 |
| **Cluster 4** | 596 | 63,3 | 345 | 36,7 |
| **Cluster 5** | 727 | 82,9 | 150 | 17,1 |
| **Cluster 6** | 114 | 14,7 | 660 | 85,3 |
| **Cluster 7** | 636 | 98,9 | 7 | 1,1 |
| **Cluster 8** | 1 | 0,1 | 703 | 99,9 |
| **Cluster 9** | 329 | 49,6 | 334 | 50,4 |
| **Cluster 10** | 377 | 63,7 | 215 | 36,3 |
| **Cluster 11** | 262 | 52,5 | 237 | 47,5 |
| **Cluster 12** | 184 | 40,8 | 267 | 59,2 |
| **Cluster 13** | 4 | 1 | 387 | 99,0 |
| **Cluster 14** | 100 | 51,8 | 93 | 48,2 |
| **Cluster 15** | 89 | 58,6 | 63 | 41,4 |
| **Cluster 16** | 44 | 55 | 36 | 45 |

**Supplementary Table 4** Primers used for RT-qPCR, qPCR and genotyping

| Gene | F primer | R primer |
|---|---|---|
| *Tet1* qPCR | GGACGCTTCGTAGCAGTACTTGAAT | CATGTACAACTGCTGTGCACCAA |
| *Tet2* qPCR | TGTTGTTGTCAGGGTGAGAATC | TCTTGCTTCTGGCAAACTTACA |
| *Tet3* qPCR | CCGGATTGAGAAGGTCATCTAC | AAGATAACAATCACGGCGTTCT |
| *Tet3* F1 genotyping | TGGATAGTATTATGCTGAGGCTCGATT | AACGCTGAGGGACTGAGGTCT |
| *Tet3* F2 genotyping | GGTCATAGCTGTGTCTGTAAACATGG | |

**Extended Table 4** Antibodies used for immunofluorescence, cell sorting and hMeDIP

| Antibody | Species | Reference | Dilution |
|---|---|---|---|
| 5-hydroxymethylcytosine | Rabbit | Active Motif, 39792 | 1:2000 |
| 5-methylcytosine | Rabbit | Active Motif, 61255 | 1:250 |
| Villin1 | Rabbit | Novus Bio, NBP1-32841 | 1:250 |
| Chga | Rabbit | Novus Bio, NBP120-15160 | 1:250 |
| Olfm4 | Rabbit | Cell signalling, #39141 | 1:100 |
| Muc2 | Rabbit | Novus Bio, NBP1-31231 | 1:250 |
| Goat anti-rabbit IgG-Alexa Fluor Plus 488 | Goat | ThermoFisher, A32731 | 1:500 |
| CD31 (PECAM-1)-PE | Rat | eBioscience, 12-0311-81 | 1:40 |
| CD45-PE | Rat | eBioscience, 12-0451-81 | 1:160 |
| TER-119-PE | Rat | eBioscience, 12-5921-81 | 1:80 |
| CD326 (EpCAM)-APC-eFluor 780 | Rat | eBioscience, 47-5791-80 | 1:160 |
